# EEG Signatures of COVID-19 Survival compared to close contacts and the Cuban EEG normative database

**DOI:** 10.1101/2024.06.21.600102

**Authors:** Ana Calzada-Reyes, Lidice Galán-García, Trinidad Virués-Alba, Lidia Charroó-Ruiz, Laura Perez-Mayo, Maria Luisa Bringas-Vega, Peng Ren, Jorge Bosh-Bayard, Yanely Acosta-Imas, Mayrim Vega-Hernández, Marlis Ontiveros-Ortega, Janet Perodin Hernandez, Eduardo Aubert-Vazquez, Deirel Paz-Linares, Joel Gutiérrez-Gil, Antonio Caballero-Moreno, Annette Valdés-Virués, Mitchell Valdés-Sosa, Roberto Rodriguez-Labrada, Pedro Valdes-Sosa

## Abstract

**Background:** The EEG constitutes a powerful neuroimaging technique for assessing functional brain impairment in COVID-19 patients.

**Objective:** The current investigation compared the EEG among COVID-19 survivors, close contacts and the Cuban EEG normative database, using semi-quantitative visual EEG inspection, quantitative and the current source density measures EEG analysis.

**Methods:** The resting-state EEG activity, quantitative QEEG, and VARETA inverse solution, were evaluated in 173 subjects: 87 patients confirmed cases by the positive reverse transcription polymerase chain reaction (RT-PCR), 86 close contacts (negative PCR) and the Cuban EEG normative database. All patients were physical, neurological, and clinically assessed using neurological retrospective survey and version 2.1 of the Schedules for Clinical Assessment in Neuropsychiatry (SCAN).

**Results:** The GTE score showed significant differences in terms of frequency scores of backgrounds rhythmic activity, diffuse slow activity, and focal abnormality. The QEEG analysis showed a pattern of abnormality with respect to the Cuban EEG normative values, displaying an excess of alpha and beta activities in the fronto-central-parietal areas in both groups. The anomalies, of COVID-19 patients and close contacts, differs in the right fronto-centro parietal area. The COVID 19 group differed-s from the close control group in theta band of the right parieto-central. The symptomatic group of COVID-19 patients differs from asymptomatic patients in delta and theta activities of the parieto-central region. The sources of activation using VARETA showed a difference in cortical activation patterns at alpha and beta frequencies in the groups studied with respect to the normative EEG database. In beta frequency were localized in right middle temporal gyrus in both groups and right angular gyrus in Covid 19 group only. In alpha band, the regions were the left supramarginal gyrus for Covid 19 group and the left superior temporal gyrus for Control group. Greater activation was found in the right middle temporal gyrus at alpha frequency in COVID-19 patients than in their close contacts.

**Conclusions:** Brain functions are impaired in long COVID-19 patients. QEEG and VARETA permit us to comprehend the susceptibility of particular brain regions exposed to viral illness.

**Highlights:**

- Background frequency abnormalities diffuse slow activity and focal abnormality associated with a pattern of excess oftheta, alpha and beta energies in in the right fronto-centro-parietal regions in QEEG analysis characterizedCOVID-19 patients.
- Patients with COVID-19 show more alpha and beta EEG activities related to normative EEG database.
- Patients with COVID-19 and close contacts show high cortical activation in temporo-parietal areas in alpha and beta bands compared to normative EEG database.
- Patients with COVID-19 (positive PCR) have high activation in the right middle frontal gyrus for alpha band related to close contacts.

## INTRODUCTION

Long-term sequelae of COVID-19 has been increasingly recognized (Desai et al., 2022;Higgins et al., 2021; Lopez-Leon et al., 2021; Taquet et al., 2021;Yong et al., 2021; Wang et al., 2020).Yet, the mechanisms of infection of the Central Nervous System (CNS) by SARS-CoV2 virus are still discussed. It has been suggested that persistence and emergence of injury neuronal, neuroinflammation, the possible viral infection of the CNS and coagulopathy could be related to neuropsychiatric long-covid symptoms (Spudich et al., 2022).

Electroencephalogram (EEG) constitutes a key neuroimaging tool for assessing the brain function (Michel et al., 2012). Also, EEG can play a significant role in the neurological brain sequelae assessment and monitoring of COVID-19 patients. Digital EEG has become progressively available in resource-limited countries. The Global Brain Consortium conference, held in 2022, launched an international call for research aim at brain dysfunctions caused by COVID-19.

Diverse studies found EEG abnormalities in COVID-19 patients (Anand et al., 2022; Chen et al., 2020; Kubota et al., 2021; Pilato et al., 2022).The most relevant EEG findings related to COVID-19 are abnormal background activity, generalized and focal slowing, epileptiform discharges with seizures, frequent sharpwave and status epilepticus, mainly described in critical COVID-19 patients (Anand et al., 2022; Chen et al., 2020; Kubota et al., 2021; Pilato et al., 2022,).Other results have reported normal EEG by visual inspection (Cecchetti et al., 2020; Helms et al., 2020; Petrescu et al., 2020). Furlanis et al., 2023 found EEG abnormalities in 65% of the sample conformed by 90 post-COVID-19 patients, prevailing a slowing activity and paroxysmal discharges mainly in the frontal regions.

Quantitative analysis of EEG (QEEG) is more sensitive than visual inspection for identifying functional brain impairment in different neuropsychiatric illness (Hughes et al., 1999; Fingelkurts et al., 2022; Livint et al., 2020). Significant differences between COVID and other causes of encephalopathies were demonstrated using QEEG (Pastor et al., 2020). A study described some QEEG findings, in critically ill COVID-19 patients, which permit predicting good neurological outcomes(Pati et al., 2020).Pastor et al., 2020 described a highly correlation between QEEG alterations and clinical evolution in a patient with an atypical delirium (Pastor et al., 2020). Increase in the delta band is associated with decreased alpha and beta bands related to control patients with a negative PCR test was reported by Saab et al., 2022. Gaber et al., 2004 found higher absolute theta power value associate with a lower level of alpha absolute power, higher level of Theta / Beta ratio and alpha and beta interhemispheric coherence abnormalities in EEG of 50 post-COVID19 patient relate to healthy individuals.

Lower delta source activity in post COVID syndrome patients in brain areas relate with executive functions was observed by Ortelly et al., 2023. Babiloni et al., 2024 recently described lower posterior resting state EEG alpha source activities in 36 post-COVID adult patients compare to normal control subjects.

Despite the accumulative information on neuropsychiatric sequelae in COVID-19 patients, very few studies have used quantitative EEG analysis to assess the abnormal findings. In this context, the aim of this research was to investigate whether alterations in the spectral power of the EEG and current density source analysis would differ between COVID-19 patients and close contact. We also examined whether the QEEG abnormalities are associated differentially with neuropsychiatric symptomatology in Cuban convalescent COVID-19 subjects.

## MATERIAL AND METHODS

### Sample

The study included 87 convalescent patients (mean age 47,14 and SD ±13,40) who were confirmed [positive reverse transcription polymerase chain reaction (RT-PCR)] cases of COVID-19 (beta variant) and monitored during 3 months after epidemiological discharge.. The control group was conformed 86 subjects (mean age 45,09 and SD ±12,65), close contacts of COVID-19 with negative RT-PCR, who were evaluated in isolation centers. The assessment was conducted, from January 2021 to December 2021, in Havana City.

### Neurological and psychiatric evaluation

The neurological assessment was performed by a trained neurologist and included physical examination and analysis of the results of the retrospective survey. The neurological survey included clinical symptomatology related to COVID-19 illness such as: cephalea, disorder of the sense of smell or taste, visual, hearing and ocular movement impairment, sensitive manifestation (superficial taste and pain), motor disorders, language, memory, conscience and sleep disturbs and dysautonomia symptomatology. For each symptom, severity and frequency of appearance were evaluated. (donde se reportan?)

The psychiatric diagnosis was achieved using the semi-structured clinical interview performed by six trained psychiatrists, the Schedules for Clinical Assessment in Neuropsychiatry (SCAN), version 2.1 (World Health Organization., WHO 1994). SCAN is a collection of instruments supported by manuals, aimed at measuring and classifying the psychopathology of the major psychiatric disorders in adult life. SCAN was developed by the WHO consisting of 1,872 items distributed in 28 sections (World Health Organization Division of Mental Health., 1994). For the present study, sections 0 (sociodemographic items); II (Physical health, somatoform and dissociative disorders) III (worries, tension etc), IV (Panic, Anxiety, and phobias); VI (Depressed mood and ideation); VII (Thinking, concentration, energy, interest); VIII (Bodily functions), X (Expansive humor and ideation) and XIII (Interference and attributions for part one) were analyzed.

### Inclusion criteria considered

(1) Subjects aged 18-80; (2) Subjects hospitalized with PCR (positive or negative); (3)3-6 months discharge period; (4) Minimum primary level of education least primary school education; and (5) absence of any current or previous history of neurological or psychiatric diseases.

### Exclusion criteria

Severe traumatic brain injury (Glasgow score of 8 or less and loss of consciousness) and severe organ-specific diseases (eg cancer, hepatopathy, cardiomyopathy and advances renal diseases) were not included.

Participation in the assessment process was voluntary; all patients and controls signed an informed consent form prior to the study.

Data of 3 patients, one with history of psychiatric illness and two children, were excluded from the study after finishing all evaluations.

### Ethics

The study was approved by the Ethics Committee of the Cuban Center for Neurosciences.

### EEG Acquisition

EEG was recorded using a 21-channel digital EEG hardware and software package from Neuronic S.A. (MEDICID 5, Cuba). Electrodes were placed in 19-electrodes sites (Fp1, Fp2, Fz, F3, F4, F7, F8, Cz, C3, C4, T3, T4, T5, T6, Pz, P3, P4, O1, and O2) according to the international 10-20 system, using surface electrodes referenced to linked earlobes with a ground electrode attached to the forehead. Impedance for all electrodes was kept less than or equal to 5 Kohm during the entire record. The EEG was amplified 1000 times, with a bandwidth of 0.5 to 30 Hz and sampled using a 12-bit analog-to-digital converter at 200 Hz. Electrodes were fitted while subjects were familiarizing with the testing equipment and procedures. EEG recording was performed in a quiet, air-conditioned room with the corresponding experimenter and recording equipment. All subjects were instructed to relax and to remain still during testing to minimize artifacts produced by ocular movements and to avoid excessive blinking. During the EEG recording, subjects were awake with their eyes closed, seated on a reclining chair. All recordings were performed during the morning. Resting EEG was obtained during 8 minutes with the eyes closed in all participants. Afterwards, it was also recorded 2 minutes of alternating close and open eyes, followed by 2 minutes of recovery.

The overall flowchart of EEG processing is displayed in Supplemental Material

### Visual EEG Analysis

Visual inspection of the EEG recordings was performed out for three Clinical neurophysiologists under blind conditions and according to the guidelines for resting state EEG analysis of the International Federation of Clinical Neurophysiology IFCN (Babiloni et al., 2020). EEG recordings were considered normal if they contained adequate organization of background activity (according to age), a well-defined spatial differentiation, rhythmic alpha activity and the absence of slow or paroxysmal wave activity. Slow EEG activity was defined as the presence of persistent non-rhythmic theta-delta slow waves. Paroxysmal activity was defined as spikes, sharp waves and polyspike-slow wave complexes. EEGs representing both types of abnormalities were included in the Slow and Paroxysmal category.

We used the global scoring scale (GTE) to quantify the group differences in the observed EEG abnormalities (De Weerd et al., 1990; Jonkman 1989). We modified the GTE score original version used in a previous study (Taboada-Crispi et al., 2018) to assess the EEG findings, accurately. More specifically, we expanded the assessment of background activity, taking into account the amplitude, and modulation of the alpha rhythm.

### Quantitative EEG analysis

We selected 20-24 EEG state closed-eyes segments (without artifact) of 2.56 s from each individual for quantitative EEG analysis by visual inspection. A minute of artifact-free EEG is considered the minimum amount of EEG required to obtain reliable quantitative measures (John et al., 1987; Nuwer et al., 1994). Fast Fourier Transform (FFT) was applied to obtain the cross spectral matrixes of all individual recordings, and were calculated with a spectral resolution of 0.39 Hz, from 0.78 to 19.53 Hz. Quantitative measures were log-transformed to obtain a normal distribution. All spectral measures were compared to gender and age-matched normative database using Z-scores.

### EEG source estimation

Variable Resolution Electromagnetic Tomography (VARETA), was used to compute, from the scalp-recorded electric potential distribution. This imaging technique allows an estimation of the distribution of the electrical generators for each frequency band within the brain, by applying a mathematical inverse solution to the EEG data. The anatomical definitions of regional probability for source localization used in VARETA are derived from a Probabilistic Brain Atlas (PBA) developed at the Montreal Neurological Institute. Resting, eyes closed EEGs from the normal population, constituted a normative database for VARETA using narrow band spectral analysis between 0.39 Hz to 19 Hz in increments of 0.39 Hz (Bosch-Bayard et al., 2001). Using the resulting set of normal values for narrow band spectral power at each scalp electrode (Valdes-Sosa et al., 1990 a,b), the sources of power at each frequency were localized. Three-dimensional color-coded tomographic images were then generated, with source generator distributions superimposed upon the transaxial, coronal, and sagittal slices color-coded as z-score deviations from the normal population.

The VARETA functional images represented the electrical activity at each voxel as squared magnitude (i.e., power) of computed current density. Current density vectors (CD) were calculated for each individual, from all the data segments, using the Neuronic Source Localizer software (Neuronic S.A.). This program provided a spatially restricted solution to cortical gray matter and basal ganglia in the Talairach Human Brain Atlas.

## Statistical analysis

### Demographic and clinical univariate analysis

Pearson’s Chi square test compared demographic, clinical and neurological variables between the two subgroups of patients studied. The level of statistical significance was set at 0.05 for all tests (Statistic 6.0 for Windows).

### Semi-quantitative EEG (sqEEG)

Using Item Response Theory (IRT), we obtained a latent variable underlying all GTE items for sqEEG. This approach is used as the indicator variables were categorical and not continuous. A polytomous IRT was implemented as the responses ranged between 0 and 5 (Beaujean, 2014; Chalmers, 2015). The analysis was conducted via R package MIRT (Chalmers, 2012), utilizing a generalized partial credit model (Pollitt and Hutchinson, 1987; Chalmers, 2015). The latent variable, “semi-quantitative Neurophysiological status” (sqNPS), was created based on the most informative items with the highest loadings. Therefore, the final latent score was the optimal combination of measured items based on their information content. Notably, the high factor loadings were solely focused on items with a clear separation of probabilities between the different levels of the item scale and not between the two groups (COVID-19 vs. Contact group), as responses for all participants were included in the IRT analysis.

### Analysis of QEEG Spectrum

The mean of EEG cross spectral parameters for both COVID-19 and close contacts were compared with the Cuban EEG normative database using the Z transform (Valdés et al., 1990). This normative EEG database was generated from the EEG of 211 normal subjects’ (105 males, 106 females) aged 5-97. Normative coefficients were obtained by carrying out a polynomial regression with age of each log spectral value. Normalized values, expressed as the number of standard deviations from the mean of the norm, were calculated for every frequency and electrode and stored as a ‘‘Z spectrum” (Valdés et al., 1990). Age might affect EEG data by increasing inter-individual variability (Szava et al., 1994). The use of normalized values for statistical analysis eliminated these effects that; otherwise, should have been taken into account for comparisons between the groups.

### Statistical methodology for comparing the Z spectra mean of both groups

A permutation test evaluated differences between the Z spectra of COVID-19 and close contacts, and Symptomatic and Asymptomatic patient groups (Blair et al., 1993; Blair et al., 1994; Galán et al., 1997; Raz., 1989). The permutation test had the following advantages: free distribution -which controls the experiment wise error for simultaneous univariate comparisons. No assumption of an underlying correlation structure, - Providing exact p-values for any number of individuals, frequency points and recording sites.

The t statistics and max (t) were calculated. Max (t) represented the maximum of t statistic in each electrode, and frequency. Multivariate statistics can be used to summarize and test differences between two Z spectra mean obtained from the maximum value of all the univariate statistics.

### Inverse solution analysis

T-test with correction for multiple comparisons identified significant regional differences in current density (CD) among COVID-19 patients, close contacts and normative EEG database for delta, theta, alpha and beta EEG bands (NeuronicStatistica software, Neuronic S.A.). The same comparison was conducted between COVID-19 patients and close contacts and between Symptomatic and Asymptomatic groups.

The outcome was a map of the t-test values for each voxel thresholder at a false discovery rate (FDR) q <0.1. Coordinates of main activation are represented in Talairach space (Neuronic Tomographic Viewer, Neuronic, SA).

## Results

### Results: Demographic and clinical variables

Analyses of demographic and clinical characteristics indicated no significant differences in age, gender and year of education between the two groups (Table 1). History of psychiatric illness, chronic obstructive pulmonary disease (CODP), renal disease, immunosuppression, cardiovascular disease and diabetes showed significant group differences. High blood pressure, asthma, neurological and other diseases did not show significant group differences (Table 1).

**Table 1.** Comparison between the COVID-19 and the close contacts of COVID-19 patient groups on demographic and clinical variables.

| Variable | Covid Group<br>n=87 | Control Group<br>n=86 | p values |
| --- | --- | --- | --- |
| **Age** | 48.85 | 44.65 | 0.42 |
| **Gender (F/M)** | 52/35 | 46/40 | 0.62 |
| **Year of Education** | 13.97 | 14.79 | 0.44 |
| **Severity of Covid** |  |  |  |
| **Asymptomatic** | 36 | - | - |
| **Mild symptomatic** | 38 | - | - |
| **Severe symptomatic** | 11 | - | - |
| **Missing** | 2 |  |  |
| **History of psychiatric illness** | 21 | 11 | 0.05* |
| **COPD** | 5 | 0 | 0.02* |
| **Renal diseases** | 6 | 0 | 0.01* |
| **Immunosuppression** | 6 | 1 | 0.05* |
| **HBP** | 33 | 22 | 0.09 |
| **Cardiovascular diseases** | 4 | 0 | 0.04* |
| **Asthma** | 10 | 10 | 0.95 |
| **Diabetes** | 7 | 1 | 0.03* |
| **Neurological disease** | 8 | 5 | 0.41 |
| **Other diseases** | 11 | 11 | 0.95 |

Pearson $\chi^2$ , $p < 0.05$ .
Gender (Female/Male F/M), COPD (Chronic obstructive pulmonary disease); History of psychiatric illness (include neurotic spectrum, neurocognitive and psychotic spectrum), HBP (high blood pressure).

Table 2 shows the neurological symptoms for COVID-19 patients and the close contacts

**Table 2.** Comparison between the COVID-19 and the close contacts of COVID-19 patient groups on neurological variables.

| Variable | Covid Group<br>n=69 | Control Group<br>n=42 | p values |
| --- | --- | --- | --- |
| **Headaches** |  |  |  |
| Headaches pre-Covid-19 | 12 | 1 | 0.67 |
| Headaches post-Covid-19 | 29 | 5 | 0.00* |
| **Taste and Smell** |  |  |  |
| Ageusia | 26 | 3 | 0.00* |
| Dysgeusia | 13 | 2 | 0.03* |
| Anosmia | 23 | 3 | 0.00* |
| Odor identification | 13 | 1 | 0.01* |
| Nasal congestion | 3 | 1 | 0.58 |
| **Hearing, Vision and Eye Movements** |  |  |  |
| Hypoacusia | 7 | 1 | 0.12 |
| Hyperacusia | 6 | 1 | 0.18 |
| Tinnitus | 4 |  | 0.40 |
| Amaurosis | 13 | 2 | 0.03* |
| Blinking lights | 9 | 1 | 0.05 |
| Visual acute | 11 | 3 | 0.17 |
| Photophobia | 6 | 2 | 0.44 |
| Blurry vision | 11 | 1 | 0.02* |
| Light adaptation | 4 | 1 | 0.39 |
| Distance adaptation | 4 | 1 | 0.39 |
| **Motor disorders** |  |  |  |
| Ataxic gait | 10 | 2 | 0.11 |
| Dizziness | 19 | 2 | 0.00* |
| Vertigo | 8 | 2 | 0.22 |
| Dysphagia | 2 | 1 | 0.87 |
| Paraparesis | 18 | 2 | 0.00* |
| Decreased muscle strength in<br>lower limbs | 9 | 2 | 0.15 |
| Precise movements | 4 | 1 | 0.41 |
| Cramps | 15 | 4 | 0.01* |
| Myalgias | 20 | 4 | 0.00* |
| Fatigue | 24 | 4 | 0.00* |
| **Sensitivity (Surface touch and Pain)** |  |  |  |
| Paresthesia | 15 | 4 | 0.09 |
| Language, Memory,<br>Consciousness and Sleep |  |  |  |
| Comprehensive aphasia | 6 | 1 | 0.18 |
| Syncope | 2 | 1 | 0.87 |
| Insomnia | 36 | 9 | 0.00* |
| Hypersomnia | 7 | 1 | 0.12 |
| **Autonomous Nervous System Disorders** |  |  |  |
| Dysuria | 5 | 1 | 0.27 |
| Diarrhea | 34 | 3 | 0.00* |
| Neurogenic syncope | 12 | 1 | 0.01* |
| Xerostomia | 24 | 1 | 0.00* |
| Hyperhidrosis | 5 | 2 | 0.59 |
| Arrhythmia | 17 | 4 | 0.04* |
| Postural hypotension | 10 | 2 | 0.11 |
| Hypertensive episodes | 9 | 3 | 0.33 |

\*Pearson $\chi^2$ , $p < 0.05$

Table 3 shows the SCAN symptoms for COVID-19 patients and the close contacts

**Table 3.**
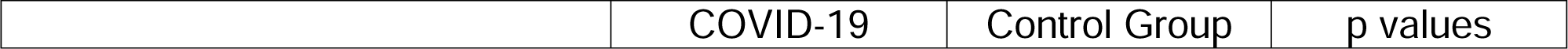
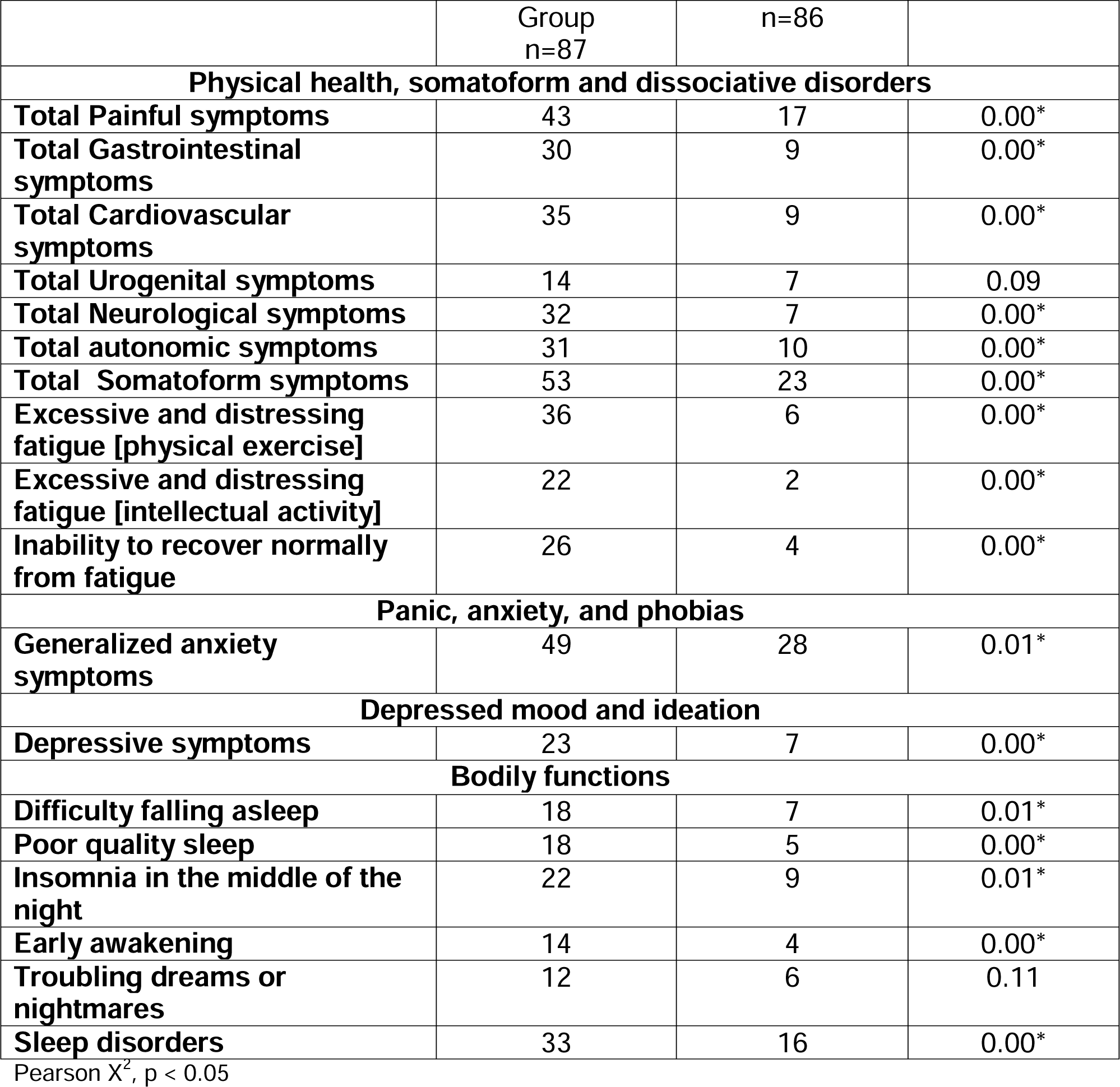
Comparison between the COVID-19 and the close contacts of COVID-19 patient groups on SCAN symptoms.

|  | COVID-19 | Control Group | p values |
| --- | --- | --- | --- |

|  | Group<br>n=87 | n=86 |  |
| --- | --- | --- | --- |
| **Physical health, somatoform and dissociative disorders** |  |  |  |
| **Total Painful symptoms** | 43 | 17 | 0.00* |
| **Total Gastrointestinal symptoms** | 30 | 9 | 0.00* |
| **Total Cardiovascular symptoms** | 35 | 9 | 0.00* |
| **Total Urogenital symptoms** | 14 | 7 | 0.09 |
| **Total Neurological symptoms** | 32 | 7 | 0.00* |
| **Total autonomic symptoms** | 31 | 10 | 0.00* |
| **Total Somatoform symptoms** | 53 | 23 | 0.00* |
| **Excessive and distressing fatigue [physical exercise]** | 36 | 6 | 0.00* |
| **Excessive and distressing fatigue [intellectual activity]** | 22 | 2 | 0.00* |
| **Inability to recover normally from fatigue** | 26 | 4 | 0.00* |
| **Panic, anxiety, and phobias** |  |  |  |
| **Generalized anxiety symptoms** | 49 | 28 | 0.01* |
| **Depressed mood and ideation** |  |  |  |
| **Depressive symptoms** | 23 | 7 | 0.00* |
| **Bodily functions** |  |  |  |
| **Difficulty falling asleep** | 18 | 7 | 0.01* |
| **Poor quality sleep** | 18 | 5 | 0.00* |
| **Insomnia in the middle of the night** | 22 | 9 | 0.01* |
| **Early awakening** | 14 | 4 | 0.00* |
| **Troubling dreams or nightmares** | 12 | 6 | 0.11 |
| **Sleep disorders** | 33 | 16 | 0.00* |

Pearson $\chi^2$ , $p < 0.05$

### Results: Visual inspection of EEG

#### Grand Total EEG Score

Analysis of the GTE using *Mirt* showed differences between groups (Z= -4.00 p-value: P(Z < z) = 0.00). However only 3 items had loadings higher than 0.7: focal abnormality, presence of sharp waves and diffuse slow activity. By contrast, the item with the lowest factor score was “Low amplitude of background activity)“.

**Table 4.** Item response theory analysis.

| ITEMS | F1 |
| --- | --- |
| 1. Frequency of rhythmic | 0.61 |
| 2. Modulation (Alpha) | 0.54 |
| 3. Low Amplitude (amplitude of background activity) | 0.11 |
| 4. Diffuse Slow Activity | 0.78** |
| 5. Reactivity BGA | 0.59 |
| 6. Paroxysmal Activity | 0.62 |
| 7. Focal Abnormalities | 0.99*** |
| 8. Sharp waves activity | 0.69* |

### Results: Quantitative EEG analysis

Significant statistical differences between the mean parameters of cross-spectral measures of COVID-19 patients and normative EEG database were found in the alpha band -a frequency range of 8.20-12.9 Hz in all EEG derivations. Within the 13.2-19.1 Hz frequency range of the beta band an increase of the energy was found at the bilateral frontal –central and parietal areas. In the band beta at within a frequency range of 13.2-Hz. The power values for these frequencies were increased in Covid-19 patients..

Significant statistical differences between the mean parameters of cross-spectral measures of close contacts and normative EEG database were found in the band alpha a frequency range of 10.9-12.9 Hz in all EEG derivations and band beta thin a frequency range of 13.2-19.1 Hz at bilateral frontal –central and right parietal area.

**Fig 1.**
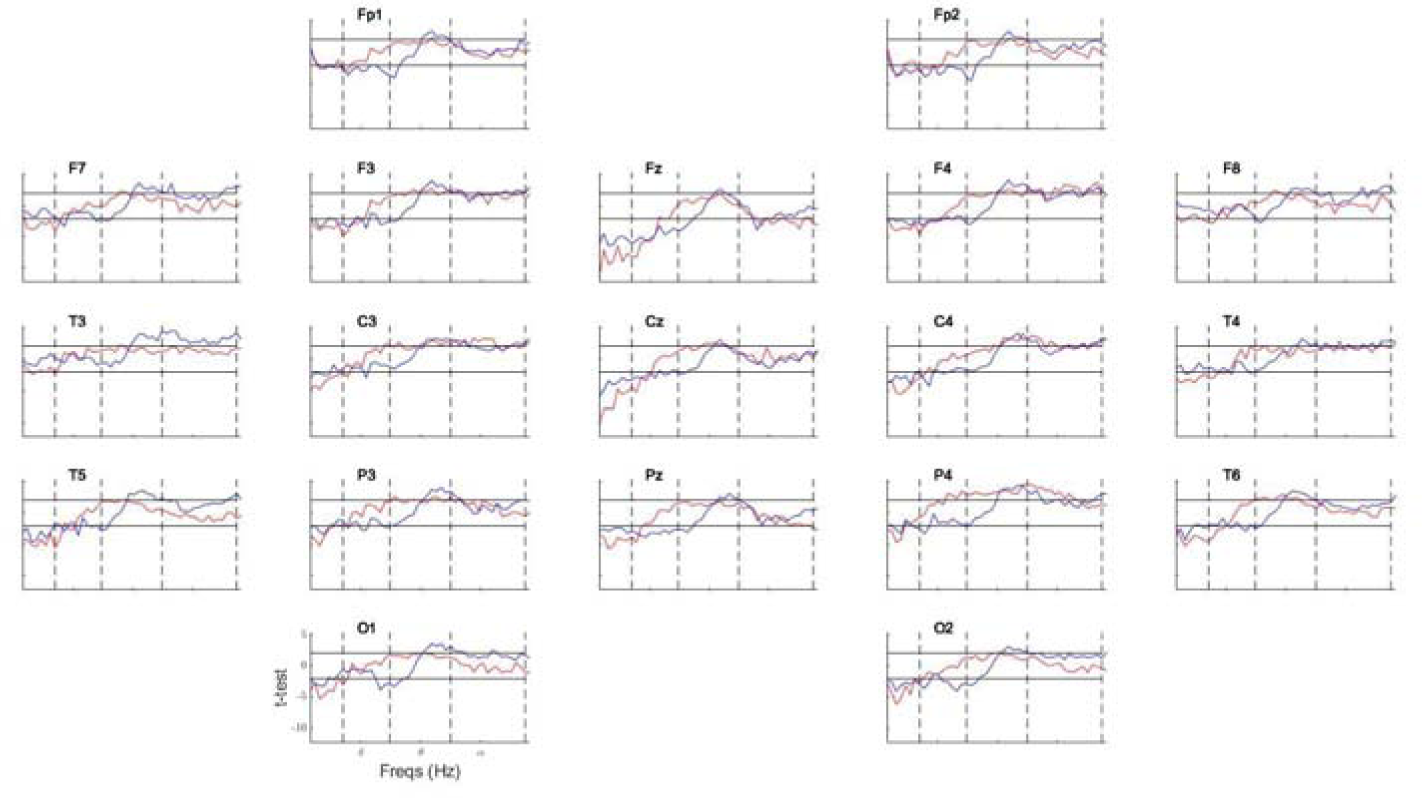
The figure shows the values of the t-Student statistic under the null hypothesis of mean against zero for each group. The red curve is for the COVID-19 group and the blue one for the Control group. The vertical dashed lines in black color mark the intervals of the classical delta, theta, alpha and beta bands respectively. The horizontal solid lines in blue color correspond to the minimum and maximum thresholds estimated using the FDR correction with q=0.10.

The permutation test allowed finding statistically significant differences for the mean of the parameters of cross-spectral measurements, in the following regions and frequencies:

### COVID-19 and Close contact groups

- Increased theta energy (frequencies ranges 5.47–5.85 Hz, 7.03-7.81 Hz) at C4, P4
– Increased alpha energy (frequencies ranges7.03-7.81 Hz) at Fp2,C4, P4
– Increased beta energy (frequency range 13.67-14.84 Hz) at C4, P4, Fz, Cz

In summary, the COVID-19 group, compared to the control, has increased energy in the theta, alpha and beta bands. The deviations are located in the right fronto- central parietal regions.

### Symptomatic and Asymptomatic groups

- Increased delta energy in 2.34 Hz at PZ
- Increased theta energy (frequencies ranges 5.85 Hz- 6.24 Hz) at Pz

In summary Symptomatic COVID-19 group when compared with the Asymptomatic groups has basically an increase of delta, theta energies at parieto- central regions.

**Fig.2.**
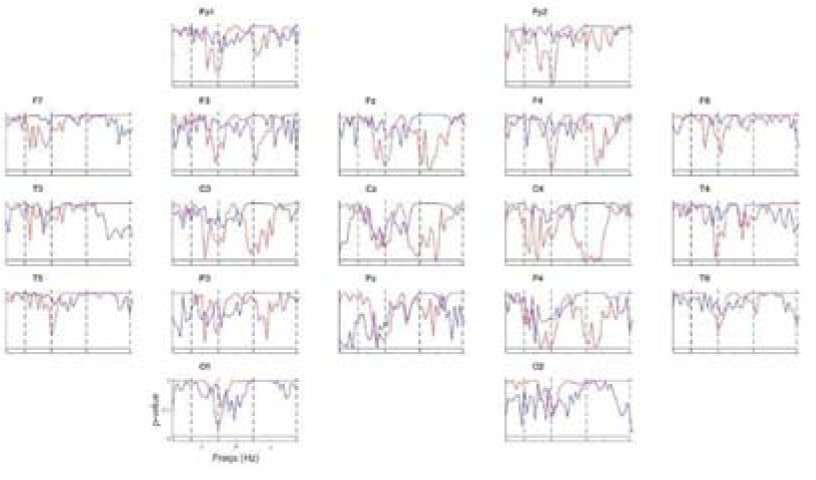
Comparison of the Z values for each frequency component (narrow band spectral frequency analysis) between patients with COVID-19 group versus close contacts of COVID-19 group (red curve) and Symptomatic and Asymptomatic patients (blue curve) by means of a permutation t-test. The scale of the abscissa axis indicates the 48 frequencies (in Hz) of the spectral analysis and the ordinate axis the p values. Each panel represents an electroencephalographic derivation. The vertical dashed lines in black color mark the intervals of the classical delta, theta, alpha and beta bands respectively. The horizontal solid lines indicate, where the probability is equal to a signification level of 0.05, significant values are below the lines.

### VARETA source analysis

VARETA source imaging revealed difference of cortical activation patterns in alpha and beta among COVID-19 group, close contacts and normative EEG database. Higher activation in alpha band located in left supramarginal area was found in COVID-19 group while in close contacts group the higher activation was in the left temporal superior gyrus (Global FDR q-Value: 0.100 after false discovery rate correction for multiple comparisons; Fig. 3).

**Fig. 3.**
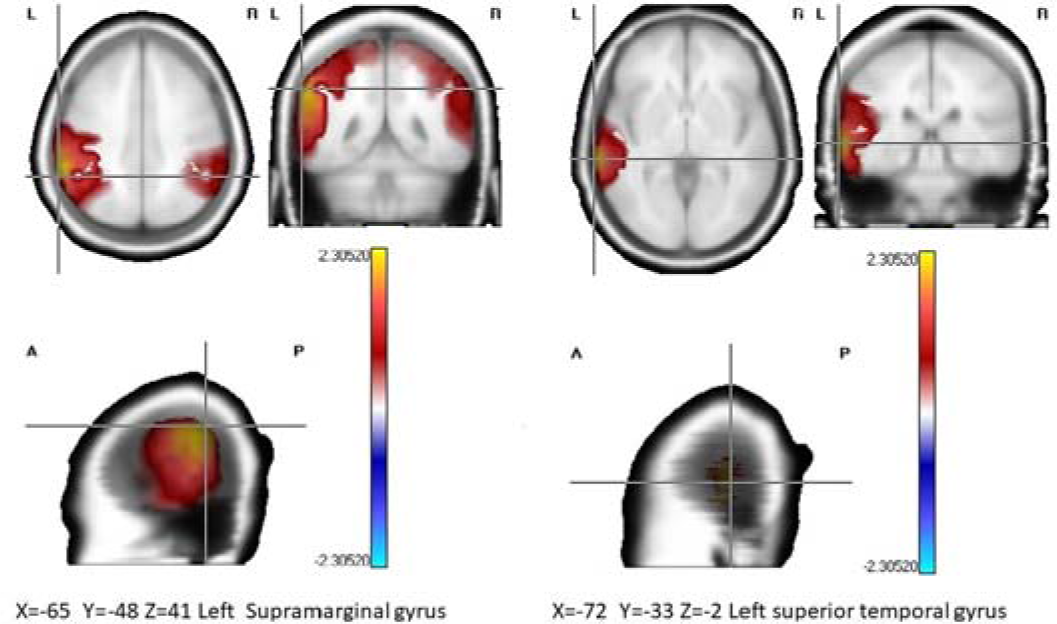
Anatomical distribution of maximum t values among patients with COVID-19, close contacts of COVID-19 patients versus normative EEG database (q-value: 0.1, after false discovery rate correction for multiple comparisons). The highest significant differences were found on the left hemisphere, mainly increase of alpha activity on the left supramarginal gyrus (COVID-19 group) and left middle temporal gyrus (close COVID-19 contact group) X, Y, Z MNI coordinates R, right hemisphere; L, left hemisphere.

Both groups revealed significant statistical differences in beta frequency related to EEG normative database at the right middle temporal gyrus and angular gyrus in COVID group.

There were no other significant group differences in any other frequency.

We found a higher activation in the right middle frontal gyrus in COVID group for alpha band related to close contacts. Global FDR q-Value: 0.100 after false discovery rate correction for multiple comparisons; Fig. 4). There were no significant differences between Symptomatic and Asymptomatic groups in any band.

**Fig. 4.**
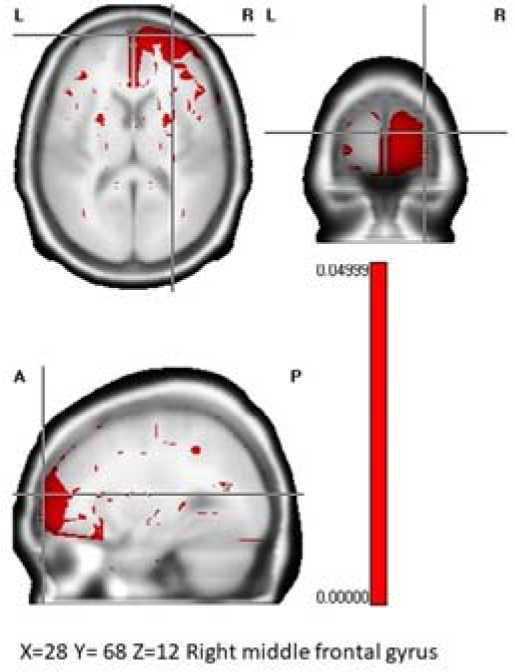
Anatomical distribution of maximum t values among patients with COVID-19 versus close contacts of COVID-19 patients (q-value: 0.1, after false discovery rate correction for multiple comparisons). The highest significant differences were found on the right hemisphere, mainly in the alpha activity on the right middle frontal gyrus (COVID-19 group) X, Y, Z MNI coordinates R, right hemisphere; L, left hemisphere.

## DISCUSSION

The effect of COVID-19 on the visual EEG, spectrum, and generating sources is a topic of ongoing research. Current studies are leading to understand better how a disease affects the brain and is reflected in the EEG. In the present study, the presence of focal abnormality diffuse slow activity and sharp wave were the most frequent abnormalities found by means of *sqEEG).* Diverse EEG studies of COVID-19 patients have reported a diffuse slowing of background in qualitative EEG analysis (Cecchettiet al., 2020; Helms et al., 2020; Ayubet al., 2021; Canhamet al., 2020; Galanopoulou et al., 2020; Louis et al., 2020; Pasini, et al.2020; Pellinen et al., 2020; Scullenet al., 2020;Sethi, 2020; Tantillo et al., 2022; Zhou et al., 2020). Regarding the effect of COVID-19 on visual EEG, a decrease in the amplitude and frequency of brain waves was observed in patients with COVID-19 compared to those without the disease (Saab et al., 2022). Recent research reported a reduction in voltage associated with nonspecific diffuse slowing EEG pattern in COVID-19 patients (Sáez-Landete et al.,2022). Another study described normal EEG or with mild diffuse slowing in COVID-19 survivors (Gogia et al., 2021). Similar abnormalities of EEG visual inspection were described for influenza A and H1N1(Kedia et al., 2011; Semmler et al., 2008). A hypothesis for explaining the EEG slowing in Covid-19 patients is the decrease of the cerebral metabolism due to the neuroinflammation caused by the viral sepsis. Other potential mechanisms are related to neuronal slowing caused by the production of pro-inflammatory molecules (Guan et al., 2020). Sharp waves mainly in frontal lobes have been reported in COVID-19 patients. (Antony & Haneef,2020, Kubota et al., 2021). This paroxysmal abnormality might be consider as a indicators of brain dysfunction caused by excitation self-sustained foci that have effects for cellular function (Zimmerman, 2013)

Both groups revealed significant statistical differences in the alpha band for all EEG derivations and in the band beta at bilateral frontal –central and parietal areas when compared to the EEG normative database. Diverse studies have described an increase of anxiety levels in confirmed case of COVID -19 and their relatives (Prakash et al., 2021, Fardin et al., 2020). Enhanced alpha activity have been described in anxious individuals (Knyazev et al., 2004). Increased levels of anxiety have been found in individuals hospitalized with a COVID-19 diagnosis (Tokur et al., 2022).

COVID-19 group had differences in theta, alpha and beta energies in the right fronto-centro-parietal regions related to the close contacts. The symptomatic group had more delta and theta activities in parieto-central regions. Regarding the effect on the EEG spectrum, studies have shown that patients with COVID-19 may present an increase in delta and theta wave activity, associated with a sleep state, and a decrease in beta wave activity related to wakefulness and concentration (Petrescu et al., 2020). Excess of theta activity could be evidence of brain dysfunction in patients with long Covid-19. Beta activity is a consequence of the thalamo/cortical and cortical/cortical interactions (Steriade, 1999). Increased beta might display brain cortical over arousal and could be relate to sleep difficulties and anxiety symptoms in patient with long Covid-19. Increase in beta EEG activity have been reported in patients with insomnia (Perlis et al., 2001; Zhao et al., 2021). Other studies in spectral analysis of COVID-19 patients, reported an encephalopathic pattern characterized by the excess of generalized delta activity and the lower alpha and beta power (Pastor et al., 2020). The main difference between our results is that Pastor et al., 2020 analyzed patients at ICU while we dealt with a few patients requiring intensive care assistance.

Our results support the hypothesis that dysfunction associated with neurotransmission contributes to the development of neuropsychiatric alterations in patients with COVID-19 (Attademo et al., 2021; Zia et al., 2022). Low oxygen levels and increased inflammatory cytokines can modify serotonin levels as a consequence of the SARS-CoV-2 virus. Low serotonin levels origin an elevated power of delta and theta activities in qEEG (Lubar, 1997) and have been clinically associated with anxiety (Østergaard, 2021). Frequent sleep impairment, reported in long COVID-19 patients, could be related to variations of serotonin and melatonin amounts (Xerfan et al., 2022). Sars-cov-2 affecting the pineal gland may reduce the action of the melatonergic pathway, thus generating circadian dysregulation and sleep disturbance, symptomatology frequent in our patients.

Moreover, our findings corroborate the theoretical hypothesis of autonomic impairment in long COVID-19 (Eshak et al., 2020).Fatigue constitutes one of the principal symptom of the dysautonomia (Barizien et al., 2021) and the most frequently reported by patients with long COVID-19 (Ceban et al., 2021). In our sample, we found excessive and distressing fatigue and inability to recover normally from fatigue in the Covid patients’ group. Other viral infections have reported similar results (Zhang et al., 2021). Increase of the delta activity have been found in patients with chronic fatigue syndrome (Zinn et al., 2018), other researchers have described increases in theta activity in the frontal, central and posterior brain regions during the mental fatigue, considering as a biomarker (Tran et al., 2020). The onset of fatigue has been associated with EEG synchronization (increase of the theta and alpha amplitude; Barwick et al., 2012; Borghini et al., 2012).

VARETA current density revealed difference in cortical activation patterns between the mean of COVID-19 and close contacts versus cero and COVID-19 group versus close contacts of COVID-19 patients. It helped to define the structures related to the increase of alpha and beta activities in COVID-19 patients and close contacts a signature of a dysfunctional neuronal state, probably secondary to central nervous system injury caused by the SARS-CoV-2 virus. Other researchers using LORETA have reported similar results to our study. Cecchettiet al (2022) found high regional current density at delta band in bilateral frontal and central-temporal regions when compared to healthy controls. A decrease in brain delta current density activity was described at rest in the frontal, parietal and temporal brain regions in 18 post COVID patients with fatigue and cognitive impairment compare to healthy subjects (Ortelly et al., 2023). Recently Babilony et al., 2024 found lower posterior resting-state electroencephalographic alpha source activities in 36 post COVID patients in comparison with normal subjects.

The possible causes of the differences found in our study could be related with the type and numbers of patients studied, the age range, and the comparison with healthy control groups. Cecchettiet al (2022) studied 49 patients recovered from COVID-19 with age range (60.8±12.6), brain aging origins loss of synaptic spines, neuronal apoptosis, and diminish in neurotransmitter levels (Hara, Rapp, Morrison, 2012). As a result, changes appear in the EEG, demonstrating how the cortical electrical delta activity diminished at the occipital regions with age (Aoki et al., 2022). Ortelly et al., (2023) included only 18 post COVID patients who’s perceiving cognitive deficits and fatigue and Babilony et al., 2024 studied post COVID patients one year after suffering from COVID 19. In the three studies the control group was integrated by healthy individuals.

In our research studied 87 post COVID patients, three to six months post-discharge, the control group was constituted by 86 close contact RT-PCR that were isolated and assessed in healthcare facilities according to the “Cuban Protocol for the Management and Care of COVID-19 in 2020” while waiting for their PCR test results. This control group was comprised to mitigate the potential influence of psychological factors, such as depression and anxiety, on intergroup differences. We use SCAN 2.0 to psychiatric assessment. Our results from the analysis of current density parameters support the hypothesis that some of the main neuropsychiatric symptomatology in patients with long COVID-19 could be associated with brain dysfunction of ACE-rich regions such as frontal, temporal and somatosensorial areas.

## Limitations

First, we performed a transversal study that simply analyzed the EEG of COVID-19 patients. Thus, it is necessary to undertake a longitudinal study to demonstrate the long Covid -19 effects in the brain. Second, we did not use a multidimensional model that included neuropsychiatric evaluation results. Consequently, future studies should consider the EEG findings within multidimensional models, comprising neuropsychiatric, behavioral and cognitive variables.

## Conclusions

Summarizing the results of this study support the hypothesis that brain functions are impaired in long COVID-19 patients. QEEG may be beneficial to understand the susceptibility of particular brain regions to viral illness and provide further insights related to the most likely electroencephalographic abnormalities in the presence of underlying neuroinflammatory mechanisms (Cantor, 2021).The combined use of semi-quantitative visual EEG inspection, the quantitative spectral EEG and VARETA increase the precision for identifying functional brain abnormalities in COVID-19 patients.

## Supporting information

Schematic representation for the methodology used for data processing

## Acknowledgments

The authors wish to thank to all EEG technicians from the Clinical Services and psychologists of the Cuban Center for Neuroscience Center, and the specialist in Psychiatry and Neurology from the Cuban Neurological and Neurosurgical Institute and other hospitals from Havana for their contribution to the assessment to the patients. We would also like to thank to the Clinical Neurophysiology Department of the Hermanos Amejeiras Hospital and the Mental Health Center in Playa municipality for their support in this research. This research was made possible by the contribution of Chengdu Science and Technology Bureau Program under Grant 2022-GH02-00042-HZ

## GRAND TOTAL SCORE

1. **Frequency of rhythmic background activity. This the predominant EEG activity.This is usually but not necessarily synonymous of alpha activity, depending of the conditions of the recordings.** 0= >9 Hz 1= 8Hz-8.9Hz 2= 7 Hz-8 Hz
2. **Amplitude of background activity** 0>31µv 1 = 16-30 2= 10-15µv
3. **Modulation of alpha rhythm** 0= Adequate modulation 1= Poor modulation 2=Without modulation
4. **Diffuse Slow activity.The presence of persistent non rhythmic theta delta slow waves localized in broader regions.** 0= None 1=Slow theta 2= intermittent theta + sporadic delta 3= intermittent theta + intermittent delta
5. **Paroxysmal activity.This is related to the activity with sudden rapid onset, rapid attainment of a maximum, and abrupt termination; distinguished from background activity, such as spikes, and spike and wave.The spikes have the duration by convention, between 20 and 70 msec.Spike and wave complex is when one spike is followed by a delta frequency wave.** 0= None 1= Paroxysmal slow activity 2=Spikes 3=Spike and wave
6. **Focal abnormality.Localization of the EEG abnormalities** 0= No focal abnormality 1= Slight unilateral abnormality 2= Slight bilateral abnormality 3=Severe unilateral + Slight contralateral 4= Severe bilateral 5= Multifocal
7. **Sharp wave activity. Paroxysmal activity that lasts from 70-200 msec** 0= None 1= Sporadic sharp waves 2=Frequent sharp waves **GTE score=(sum 1-7)+1**

## REFERENCES

1. Desai AD, Lavelle M, Boursiquot BC, Wan EY. Long-term complications of COVID-19.Am J Physiol Cell Physiol.2022;322(1):C1–C11. doi: 10.1152/ajpcell.00375.2021

2. Higgins V, Sohaei D, Diamandis EP, Prassas I. COVID-19: from an acute to chronic disease?Potential long-term health consequences.Crit Rev Clin Lab Sci. 2021;58(5):297–310.doi: 10.1080/10408363.2020.1860895.

3. Lopez-Leon S, Wegman-Ostrosky T, Perelman C, Sepulveda R, Rebolledo PA, Cuapio A, et al. More than 50 long-term effects of COVID-19: a systematic review and meta-analysis. Sci Rep. 2021; 11(1):16144. doi: 10.1038/s41598-021-95565-8

4. Taquet M, Dercon Q, Luciano S, Geddes JR, Husain M, Harrison PJ. Incidence, co-occurrence, and evolution of long-COVID features: A 6-month retrospective cohort study of 273,618 survivors of COVID-19.PLoS Med. 2021;18(9):e1003773. doi: 10.1371

5. Yong SJ.Long COVID or post-COVID-19 syndrome: putative pathophysiology, risk factors, and treatments.Infect Dis (Lond).2021;53(10):737–754. doi: 10.1080/23744235.2021.1924397.

6. Wang F, Kream RM, Stefano GB. Long-Term Respiratory and Neurological Sequelae of COVID-19.Med SciMonit. 2020 Nov 1;26:e928996. doi: 10.12659/MSM.928996

7. Spudich S, Nath A. Nervous system consequences of COVID-19.Science.2022;375(6578):267-269. doi: 10.1126/science.abm2052.

8. Michel CM, Murray MM.Towards the utilization of EEG as a brain imaging tool.Neuroimage.2012; 61(2):371–85. doi:10.1016/j.neuroimage.2011.12.039

9. Valdes-Sosa PA, Evans AC, Valdes-Sosa MJ, Poo MM. A call for international research on COVID-19-induced brain dysfunctions. Natl Sci Rev. 2021 Oct 27;8(12):nwab190. doi: 10.1093/nsr/nwab190.

10. Anand P, Al-Faraj A, Sader E, Dashkoff J, Abdennadher M, Murugesan R, et al. A. Seizure as the presenting symptom of COVID-19: A retrospective case series.Epilepsy Behav.2020;112:107335. doi: 10.1016/j.yebeh.2020.107335.

11. Chen W, Toprani S, Werbaneth K, Falco-Walter J. Status epilepticus and other EEG findings in patients with COVID-19: a case series.Seizure.2020;81:198–200.

12. Kubota T, Prasannakumar KG, Naoto K. Meta-analysis of EEG findings in patients with COVID-19.Epilepsy Behav.2021;115:107682. 10.1016/j.yebeh.2020.107682.

13. Pilato MS, Urban A, Alkawadri R, Barot NV, Castellano JF, Rajasekaran V, et al. EEG Findings in Coronavirus Disease. J Clin Neurophysiol. 2022; 39(2):159–165. doi: 10.1097/WNP.0000000000000752.

14. Cecchetti G, Vabanesi M, Chiefo R, Fanelli G, Minicucci F, Agosta F, et al. Cerebral involvement in COVID-19 is associated with metabolic and coagulation derangements: an EEG study. J Neurol 2020:267:3130–3134

15. Helms J, Kremer S, Merdji H, Clere-Jehl R, Schenck M, Kummerlen C, et al. Neurologic Features in severe SARS-CoV-2 infection. N Engl J Med. 2020;382(23):2268–2270. doi: 10.1056/NEJMc2008597.

16. Petrescu AM, Taussig D, Bouilleret V. Electroencephalogram (EEG) in COVID-19: A systematic retrospective study.NeurophysiolClin. 2020 Jul;50(3):155–165. doi: 10.1016/j.neucli.2020.06.001.

17. Furlanis G, Buoite Stella A, Biaduzzini F, Bellavita G, Frezza NA, Olivo S, et al. Cognitive deficit in post-acute COVID-19: an opportunity for EEG evaluation?. Neurol Sci. 2023;44(5):1491–8. doi: 10.1007/s10072-023-06615-0

18. Hughes JR, John ER. Conventional and quantitative EEG. J Neuopsych UinNeurosci1999; 11:190.208.

19. Fingelkurts AA, Fingelkurts AA. Quantitative Electroencephalogram (qEEG) as a natural and non-Invasive window into living brain and mind in the functional continuum of healthy and pathological conditions. Applied Sciences. 2022;12(19):9560.

20. Livint Popa L, Dragos H, Pantelemon C, VerisezanRosu O, Strilciuc S. The Role of Quantitative EEG in the Diagnosis of Neuropsychiatric Disorders.J Med Life.13 (1):8–15. doi: 10.25122/jml-2019-0085

21. Pastor J, Vega-Zelaya L, Martín Abad E. Specific EEG encephalopathy pattern in SARS-CoV-2 Patients.J Clin Med. 2020;9(5):1545.doi: 10.3390/jcm9051545.

22. Pati S, Toth E, Chaitanya G. Quantitative EEG markers to prognosticate critically ill patients with COVID-19: a retrospective cohort study. Clin Neurophysiology.2020;131(8):1824.

23. Saab C, Valsamis H, Baki S, Leung J, Ghosn S, Lapin B, et al. SARS-CoV-2 slows brain rhythms with more severe effects in younger individuals.2022; doi;/10.21203/rs.3.rs-1197196/v1.

24. Gaber MM, Hosny H, Hussein M, Ashmawy MA, Magdy R. Cognitive function and quantitative electroencephalogram analysis in subjects recovered from COVID-19 infection. BMC Neurol. 2024;24(1):60. doi: 10.1186/s12883-023-03518-7.

25. Ortelli P, Quercia A, Cerasa A, Dezi S, Ferrazzoli D, Sebastianelli L, Saltuari L, Versace V, Quartarone A. Lowered Delta Activity in Post-COVID-19 Patients with Fatigue and Cognitive Impairment. Biomedicines. 2023;11(8):2228. doi: 10.3390/biomedicines11082228.

26. Babiloni C, Cacciola EG, Tucci F, Vassalini P, Chilovi A, Jakhar D, et al. Resting-state EEG rhythms are abnormal in post COVID-19 patients with brain fog without cognitive and affective disorders. Clinical Neurophysiology. 2024;161:159–72. doi:10.1016/j.clinph.2024.02.034

27. World Health Organization. Schedules for clinical assessment in neuropsychiatry, version 2.1.Geneva: World Health Organization.1994.

28. Babiloni C, Barry RJ, Başar E, Blinowska KJ, Cichocki A, Drinkenburg WH, et al. International Federation of Clinical Neurophysiology (IFCN)–EEG research workgroup: Recommendations on frequency and topographic analysis of resting state EEG rhythms. Part 1: Applications in clinical research studies. Clinical Neurophysiology.2020;131(1):285-307.

29. De Weerd A, Perquin W, Jonkman E. Role of the EEG in the prediction of dementia in Parkinson’s disease.Dementia.1990; 1: 115–118. doi: 10.1159/000107129

30. Jonkman EA.Simple EEG scoring method for senile dementia of the Alzheimer type.Electroenceph. Clin. Neurophysiol.1989.72:44.

31. Taboada-Crispi A, Bringas-Vega ML, Bosch-Bayard J, Galán-García L, Bryce C, Rabinowitz AG, et al. Quantitative EEG tomography of early childhood malnutrition.Front Neurosci.2018;12:595.

32. John ER, Prichep LS, Easton P. Normative data banks and neurometrics: basic concepts, methods and results of norm construction.In: Remond A, editor.Handbook of electroencephalography and clinical neurophysiology.Computer analysis of the EEG and other neurophysiological signals, vol. III. Amsterdam: Elsevier; 1987.p. 449-95.

33. Nuwer MR, Lehmann D, Lopes da Silva F, Matsuoka S, Sutherling W Vibert JF.IFCN guidelines for topographic and frequency analysis of EEGs and EPs. Electroencephalogr Clin Neurophysiol 1994;91:1–5 John ER, Karmel BZ, Corning WC, Easton P, Brown D, Ahn H, et al. Neurometrics: Numerical taxonomy identifies different profiles of brain functions within groups of behaviorally similar people. Science. 1977; 196: 1393-1410.

34. Bosch-Bayard, J, Valdés-Sosa, P., Virues-Alba, T., Aubert-Vázquez, E., John, E. R., Harmony, T., et al. (2001). 3D statistical parametric mapping of EEG source spectra by means of variable resolution electromagnetic tomography (VARETA). Clin. EEG 32, 47–61. doi: 10.1177/155005940103200203

35. Valdes-Sosa P, Bosch J, Gray F, Hernandez J, Riera J, Pascual R, et al. Frequency domain models of the EEG. Brain Topog. 1992; 4: 309–319.

36. Valdes-Sosa P, Biscay R, Galan L, Bosch J, Szava S, Virues T. High resolution spectral EEG norms for topography. Brain Topography. 1990a; 3: 281–282.

37. Valdés PA, Biscay R, Galán L, Bosch J, Száva S, Virués T. High resolution spectral norms for topography.Brain Topogr 1990b;32:281–2.

38. Beaujean A A. Latent Variable Modeling Using R: A Step-by-Step Guide (1st ed.). Routledge: New York. 2014.doi: 10.4324/9781315869780

39. Chalmers RP. Extended mixed-effects item response models with the MH-RM algorithm.. J. Educ. Meas. 2015;52(2):200–22.

40. Chalmers RP. mirt: A multidimensional item response theory package for the R environment. J. Stat. Softw. 2012;48:1-29.

41. Pollitt E, Gorman KS, Engle PL, Martorell R, Rivera J. Early supplementary feeding and cognition: effects over two decades. Monogr Soc Res Child Dev. 1993;58(7):1–99; discussion 111-8. doi: 10.2307/1166162.

42. Szava S, Valdés P, Biscay R, Galán L, Bosch J, Clark I, et al. High resolution quantitative EEG analysis.Brain Topogr 1994;6:211–9.

43. Blair R, Karniski RW. An alternative method for significance testing of waveform difference potential. Psychophysiology 1993;30:518–24.

44. Blair R, Karniski W. Distribution-free statistical analyses of surface and volumetric maps. In: Thatcher RW, Hallett M, Zeffiro T, John ER, Huerta M, editors. Functional neuroimaging. New York: Academic Press; 1994. p. 19-28.

45. Galán L, Biscay R, Rodríguez JL, Pérez Abalo MC, Rodríguez R. Testing topographic differences between events related brain potentials by using nonparametric combinations of permutation test. ElectroencephalogrClinNeurophysiol 1997;102:240–7.

46. Raz J. Analysis of repeated measurements using non-parametric smoothers and randomization tests.Biometrics 1989;45:851–71.

47. Ayub N, Cohen J, Jing J, Jain A, Tesh R, Mukerji SS, Zafar SF, Westover MB, Kimchi EY.Clinical Electroencephalography findings and considerations in hospitalized patients With Coronavirus SARS-CoV-2. Neurohospitalist. 2021;11(3):204–213. doi: 10.1177/1941874420972237.

48. Canham LJW, Staniaszek LE, Mortimer AM, Nouri LF, Kane NM. Electroencephalographic (EEG) features of Covid-19: a case series.ClinNeurophysiolPract. 2020;5:199–205. doi:10.1016/j.cnp.

49. Galanopoulou AS, Ferastraoaru V, Correa DJ, Cherian K, Duberstein S, Gursky J, et al. EEG fndings in acutely ill patients investigated for SARS-CoV-2/COVID-19: A small case series preliminary report.Epilepsia Open.2020;2:314–324.

50. Louis S, Dhawan A, Newey C, Nair D, Jehi L, Hantus S, et al. Continuous electroencephalography characteristics and acute symptomatic seizures in COVID-19 patients.ClinNeurophysiol. 2020;131(11):2651–2656. doi: 10.1016/j.clinph.2020.08.003

51. Pasini E, Bisulli F, Volpi L, Minardi I, Tappatà M, Muccioli L, et al. EEG findings in COVID-19 related encephalopathy.ClinNeurophysiol. 2020;131(9):2265–2267. doi: 10.1016/j.clinph.2020.07.003

52. Pellinen, J, Carroll, E, Friedman, D, Boffa M, Dugan P, Friedman DE, et al. Continuous EEG findings in patients with COVID-19 infection admitted to a New York academic hospital system.Epilepsia.2020;61(10):2097–105. 10.1111/epi.16667.

53. Scullen T, Keen J, Mathkour M, Dumont AS, Kahn L. Coronavirus 2019 (COVID-19)-Associated Encephalopathies and Cerebrovascular Disease: The New Orleans Experience.World Neurosurg. 2020;141:e437–e446. doi: 10.1016/j.wneu.2020.05.192

54. Sethi NK.EEG during the COVID-19 pandemic: What remains the same and what is different.ClinNeurophysiol. 2020;131(7):1462. doi: 10.1016/j.clinph.2020.04.007

55. Vespignani H, Colas D, Lavin BS, Soufflet C, Maillard L, PourcherV,et al.. Report on Electroencephalographic Findings in Critically Ill Patients with COVID-19. Ann Neurol. 2020;88(3):626–630. doi: 10.1002/ana.25814.

56. Tantillo GB, Jette N, Gururangan K, Agarwal P, Marcuse L, Singh A, Goldstein J, Kwon CS, Dhamoon MS, Navis A, Nadkarni GN.Electroencephalography at the height of a pandemic: EEG findings in patients with COVID-19.ClinNeurophysiol. 2022;137:102–12.

57. Zhou Y, Guan J, Li Y, Li H., Li X. EEG patterns in patients with COVID-19: A preliminary study.Brain and behavior.2020. 10(10): e01905

58. Sáez-Landete I, Gómez-Domínguez A, Estrella-León B, Díaz-Cid A, Fedirchyk O, Escribano-Muñoz M, et al. Retrospective Analysis of EEG in Patients with COVID-19: EEG Recording in Acute and Follow-up Phases. Clin EEG Neurosci. 2022;53(3):215–228. doi: 10.1177/15500594211035923.

59. Gogia B, Thottempudi N, Ajam Y, Singh A, Ghanayem T, Dabi A, Fang X, Masel T, Rai P. EEG characteristics in COVID-19 survivors and non-survivors with seizures and encephalopathy.Cureus. 2021;13(10).

60. Ekstrand JJ, Herbener A, Rawlings J, Turney B, Ampofo K, Korgenski EK, Bonkowsky JL. Heightened neurologic complications in children with pandemic H1N1 influenza. Ann Neurol. 2010;68(5):762–6. doi: 10.1002/ana.22184

61. Kedia S, Stroud B, Parsons J, Schreiner T, Curtis DJ, Bagdure D, et al. Pediatric neurological complications of 2009 pandemic influenza A (H1N1). Arch Neurol. 2011;68(4):455–62. doi: 10.1001/archneurol.2010.318

62. Semmler A, Hermann S, Mormann F, Weberpals M, Paxian SA, Okulla T, et al. Sepsis causes neuroinflammation and concomitant decrease of cerebral metabolism. J Neuroinflammation. 2008;5:38. doi: 10.1186/1742-2094-5-38.

63. Guan J, Li Y, Li H, Li X, Liu X. EEG features of patients with COVID-19: A preliminary study.Brain and behavior.2020: e01705

64. Antony AR, Haneef Z. Systematic review of EEG findings in 617 patients diagnosed with COVID-19. Seizure. 2020;83:234–41

65. Zimmerman EM. Focal Sharp Waves in Psychiatric Patients: Implications for Complex Clinical Presentation. The Chicago School of Professional Psychology; 2013.

66. Prakash J, Dangi A, Chaterjee K, Yadav P, Srivastava K, Chauhan VS. Assessment of depression, anxiety and stress in COVID-19 infected individuals and their families. Med J Armed Forces India. 2021;77(Suppl 2):S424–S429. doi: 10.1016/j.mjafi.2021

67. Fardin MA. COVID-19 and anxiety: a review of psychological impacts of infectious disease outbreaks. Arch Clin Infect Dis. 2020; 15(COVID-19).10.5812/archcid.102779

68. Knyazev GG, Savostyanov AN, Levin EA. Alpha oscillations as a correlate of trait anxiety. Int J Psychophysiol. 2004;53(2):147–60. doi: 10.1016/j.ijpsycho.2004.03.001

69. TokurKesgin M, HançerTok H, Uzun LN, Pehlivan Ş. Comparison of anxiety levels of hospitalized COVID-19 patients, individuals under quarantine, and individuals in society. PerspectPsychiatr Care. 2022; 58(1):149–158. doi: 10.1111/ppc.12857

70. Steriade M. Cellular substrates of brain rhythms. In: Niedermeyer E, Lopes da Silva FH, editors. Electroencephalography e basic principles, clinical applications, and related fields.4th ed. Baltimore: Williams &Wilkins; 1999.p. 28-75.

71. Perlis ML, Merica H, Smith MT, Giles DE.Beta EEG activity and insomnia.Sleep Med Rev. 2001;5(5):365–76.

72. Zhao W, Van Someren EJ, Li C, Chen X, Gui W, Tian Y, Liu Y, Lei X. EEG spectral analysis in insomnia disorder: A systematic review and meta-analysis.Sleep Medicine Reviews.2021 Oct 1;59:101457.

73. Attademo L, Bernardini F. Are dopamine and serotonin involved in COVID-19 pathophysiology? Eur J Psychiatry. 2021;35(1):62–63. doi: 10.1016/j.ejpsy.2020.10.004.

74. Zia N, Ravanfar P, Allahdadian S, Ghasemi M. Impact of COVID-19 on neuropsychiatric disorders.J Clin Med. 2022 Sep 3;11(17):5213. doi: 10.3390/jcm11175213

75. Lubar JF.Neocortical dynamics: implications for understanding the role of neurofeedback and related techniques for the enhancement of attention.ApplPsychophysiol Biofeedback.1997 Jun;22(2):111–26. doi: 10.1023/a:1026276228832

76. Østergaard L. SARS CoV-2 related microvascular damage and symptoms during and after COVID-19: Consequences of capillary transit-time changes, tissue hypoxia and inflammation.Physiol Rep. 2021 Feb;9(3):e14726.doi: 10.14814/phy2.14726.

77. Xerfan EMS, Morelhao PK, Arakaki FH, Facina ADS, Tomimori J, Xavier SD, Tufik S, Andersen ML.Could melatonin have a potential adjuvant role in the treatment of the lasting anosmia associated with COVID-19?A review.Int J Dev Neurosci.2022 Oct;82(6):465–470. doi: 10.1002/jdn.10208.

78. Eshak N, Abdelnabi M, Ball S, Elgwairi E, Creed K, Test V, Nugent K. Dysautonomia: An overlooked neurological manifestation in a critically ill COVID-19 Patient. Am J Med Sci. 2020;360(4):427–429. doi: 10.1016/j.amjms.2020.07.022.

79. Barizien N, Le Guen M, Russel S, Touche P, Huang F, Vallée A. Clinical characterization of dysautonomia in long COVID-19 patients. Sci Rep. 2021;11(1):14042. doi: 10.1038/s41598-021-93546-5.

80. Ceban F, Ling S, Lui LMW, Lee Y, Gill H, Teopiz KM, et al. Fatigue and cognitive impairment in Post-COVID-19 Syndrome: A systematic review and meta-analysis.Brain Behav Immun. 2021;101: 93-135

81. Zhang X, Wang F, Shen Y, Zhang X, Cen Y, Wang B, et al. Symptoms and health outcomes among survivors of COVID-19 infection 1 Year after discharge from hospitals in Wuhan, China. JAMA Netw Open.2021; 4(9):e2127403. doi: 10.1001/jamanetworkopen.2021.27403.

82. Zinn MA, Zinn ML, Valencia I, Jason LA, Montoya JG. Cortical hypoactivation during resting EEG suggests central nervous system pathology in patients with chronic fatigue syndrome.Biol Psychol.2018;136:87–99.doi: 10.1016/j.biopsycho.2018.05.016.

83. Tran Y, Craig A, Craig R, Chai R, Nguyen H.The influence of mental fatigueon brainactivity: Evidence from a systematicreviewwithmeta-analyses.Psychophysiology.2020; 57(5):e13554. doi: 10.1111/psyp.13554.

84. Barwick F, Arnett P, Slobounov S. EEG correlates of fatigue during administration of a neuropsychological test battery.Clin Neurophysiol. 2012;123(2):278–84.

85. Borghini G, Vecchiato G, Toppi J, Astolfi L, Maglione A, Isabella R, et al. Assessment of mental fatigue during car driving by using high resolution EEG activity and neurophysiologic indices. Annu Int Conf IEEE Eng Med Biol Soc.2012;6442–5.doi: 10.1109/EMBC.2012.6347469.

86. Hara Y, Rapp PR, Morrison JH. Neuronal and morphological bases of cognitive decline in aged rhesus monkeys. Age. 2012;34:1051–73.doi: 10.1007/s11357-011-9278-5

87. Aoki Y, Hata M, Iwase M, Ishii R, Pascual-Marqui RD, Yanagisawa T, et al. Cortical electrical activity changes in healthy aging using EEG-eLORETA analysis. Neuroimage: Reports, 2(4):100143. doi: /10.1016/j.ynirp.2022.100143

88. Cantor D. COVID-19: Effects on brain and qEEG correlates-a window into the Future. Int J Psychophysiol. 2021; 168:S83. doi: 10.1016/j.ijpsycho.2021.07.257.

