## Supplementary material for "EEG Signatures of COVID-19 Survival compared to close contacts and the Cuban EEG normative database": Schematic representation for the methodology used for data processing

The methodology for obtaining and processing data is summarized in the following steps:

1- Acquisition of measurements derived from the application of neurological and psychiatric batteries and EEG recording

2- Extraction of descriptive parameters (DPs)

3- Statistical analyzes to contrast the differences between the Covid group and the close contact group.

**Step 1: Acquisition of measurements**

*Neurological and psychiatric evaluation*

The neurological assessment was performed by a trained neurologist and included physical examination and analysis of the results of the retrospective survey. The neurological survey included clinical symptomatology related to COVID-19 illness such as: cephalea, disorder of the~~ir~~ sense of smell or taste, visual, hear and ocular movement impairment, sensitive manifestation (superficial taste and pain), motor disorders, language, memory, conscience and sleep disturbs and dysautonomia symptomatology. For each symptom, severity and frequency of appearance were evaluated.

The psychiatric diagnosis was made using the semi-structured clinical interview performed by a six trained psychiatrists, the Schedules for Clinical Assessment in Neuropsychiatry (SCAN), version 2.1 (World Health Organization., 1994). SCAN is a collection of instruments supported by manuals, aimed at measuring and classifying the psychopathology of the major psychiatric disorders ~~of~~ in adult life. SCAN was developed by the World Health Organization and consists of 1,872 items distributed in 28 sections (World Health Organization Division of Mental Health., 1994). For the present study, sections 0 (sociodemographic items); II (Physical health, somatoform and dissociative disorders) III (worries, tension etc), IV (Panic, Anxiety, and phobias); VI (Depressed mood and ideation); VII (Thinking, concentration, energy, interest); VIII (Bodily functions), X (Expansive humor and ideation) and XIII (Interference and attributions for part one) were analyzed.

*EEG Acquisition*

EEG was recorded using a 21-channel digital EEG hardware and software package from Neuronic S.A. (MEDICID 5, Cuba). Electrodes were placed in 19-electrodes sites (Fp1, Fp2, Fz, F3, F4, F7, F8, Cz, C3, C4, T3, T4, T5, T6, Pz, P3, P4, O1, and O2) according to the international 10-20 system, using surface electrodes referenced to linked earlobes with a ground electrode attached to the forehead. Impedance for all electrodes was kept less than or equal to 5 Kohm during the entire record. The EEG was amplified 1000 times, with a bandwidth of 0.5 to 30 Hz and sampled using a 12-bit analog-to-digital converter at 200 Hz. Electrodes were fitted while subjects were familiarizing with the testing equipment and procedures. EEG record was performed in a quiet, air-conditioned room with the corresponding experimenter and recording equipment. All subjects were instructed to relax and to remain still during testing to minimize artifacts produced by ocular movements and to avoid excessive blinking. In recording, subjects were awake with the eyes closed, seated on a reclining chair. All recordings were performed during the morning. Resting EEG was obtained during 8 minutes with the eyes closed in all participants. Afterwards, it was also recorded 2 minutes of alternating close and open eyes, followed by 2 minutes of recovery.

**Step 2: Extracted descriptive parameters (DPs)**

*Neurological Battery and Psychiatric Battery:*

The presence of appearance and severity was determined for each symptom.

*EEG:*

The following parameters were extracted *from the EEG:*

*The global scoring scale (GTE)*

- The presence of Frequency of rhythmic background activity, Modulation, Low amplitude, Diffuse slow activity, Reactivity, Paroxysmal activity, Focal abnormality and Sharp wave were evaluated.
- *The cross spectral matrixes*
- *The ESI parameter*

Figures 1 and 2 show the workflow for the extraction of the spectrum parameters, cross spectral matrix and source generators.

The Fast Fourier Transform (known as FFT) was applied to those artifact-free EEG epochs to find the complex EEG amplitude spectra in the frequency domain. The absolute value of the source cross-spectra, estimated as the covariances of complex solutions in all frequencies. These values were the input to calculate the Electrophysiological Source Imaging (ESI) for all 49 frequencies from 0.39 to 19.14 Hz with step of 0.39 Hz (Bosch-Bayard et al., 2001).

The EEG source activity and connectivity with Electrophysiological Source Imaging (ESI) were estimated using BC-Vareta method ((Bosch-Bayard et al., 2001). The method is considered within the flexible Multiple Penalized Least Squares (MPLS) framework, which allows a common formulation for combine different degree of smoothness and sparseness as restrictions or contrasts. The solution can be found from the optimization problem:

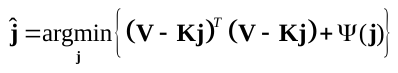

where the penalty term takes the form of a sum of several constraints or penalty functions, i.e.,
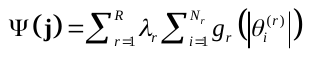
. This is evaluated at the components of the vector
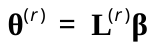
, with
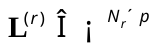
 being linear operators that impose a structural relationship among coefficients.

A standard template head model was used for estimating ESI on all subjects. The generating sources were defined over a cortical surface mesh containing 5656 vertices on brain regions that are physiologically feasible EEG generators (e.g., avoiding vertices in corpus callosum). Using a three-sphere piece-wise homogenous and isotropic head model, the projection matrix (i.e., the Electric Lead Field) was computed for the standard positions of the array of 19 electrodes from the 10/20 system in Neuronic Source Localizer (Riera & Fuentes, 1998).

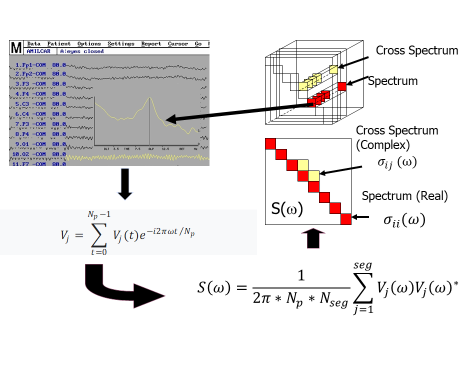
.

Figure 1: Workflow for extraction of the spectrum and cross spectrum matrix

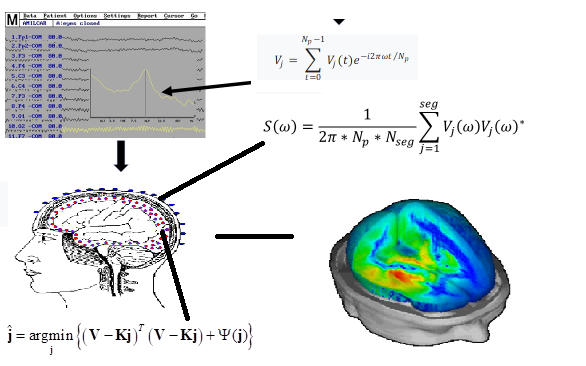

Figure 2: Workflow for extraction of the ESI parameters

**Step 3- Statistical analyzes to contrast the differences between the Covid group and the close contact group.**

- Comparison of DPs with respect to a normative database. Univariate “z-transformation” of DPs to a metric defined by the normative database.
- Comparison between vectors of means using permutation techniques

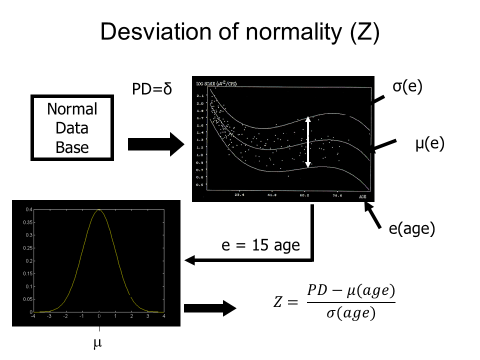

Figure 3: Workflow for comparison with database normative,

where μ(age) and σ (age) are parameters which describe a polynomial regression with respect to the age in the population
